# Quantifying the effects of switchgrass (*Panicum virgatum*) on deep organic C stocks using natural abundance ^14^C in three marginal soils

**DOI:** 10.1101/2020.07.07.190587

**Authors:** Eric W. Slessarev, Erin E. Nuccio, Karis J. McFarlane, Christina Ramon, Malay Saha, Mary K. Firestone, Jennifer Pett-Ridge

## Abstract

Perennial bioenergy crops have been shown to increase soil organic carbon (SOC) stocks, potentially offsetting anthropogenic C emissions. The effects of perennial bioenergy crops on SOC are typically assessed at shallow depths (< 30 cm), but the deep root systems of these crops may also have substantial effects on SOC stocks at greater depths. We hypothesized that deep (> 30 cm) soil organic carbon (SOC) stocks would be greater under bioenergy crops relative to stocks under shallow-rooted conventional crop cover. To test this, we sampled soils to between 1- and 3-meters depth at three sites in Oklahoma with 10-20 year old switchgrass (*Panicum virgatum*) stands, and collected paired samples from nearby fields cultivated with shallow rooted annual crops. We measured root biomass, total organic C, ^14^C, ^13^C, and other soil properties in three replicate soil cores in each field and used a mixing model to estimate the proportion of recently fixed C under switchgrass based on ^14^C. The subsoil C stock under switchgrass (defined over 500-1500 kg m^-2^ equivalent soil mass, approximately 30-100 cm depth) exceeded the subsoil stock in neighboring fields by 1.5 kg C m^-2^ at a sandy loam site, 0.6 kg C m^-2^ at a site with loam soils, and showed no significant difference at a third site with clay soils. Using the mixing model, we estimated that additional SOC introduced after switchgrass cultivation comprised 31% of the subsoil C stock at the sandy loam site, 22% at the loam site, and 0% at the clay site. These results suggest that switchgrass can contribute significantly to subsoil organic C—but also indicated that this effect varies across sites. Our analysis shows that agricultural strategies that emphasize deep-rooted grass cultivars can increase soil C relative to conventional crops while expanding energy biomass production on marginal lands.

## 1. Introduction

Soil horizons deeper than 30 cm contain the majority of Earth’s soil organic carbon (SOC)— possibly holding well over 1000 Pg of C globally (Harrison et al., 2010; Jobbágy and Jackson, 2000). While the bulk of deep soil C tends to exchange slowly with the atmosphere (Mathieu et al., 2015; Trumbore, 2009), SOC losses from deep soil horizons following land use change have been substantial—accounting for the majority of the 133 Pg of SOC lost following the global expansion of agriculture (Sanderman et al., 2017). By implication, successful attempts to reverse SOC loss in agricultural lands must restore SOC in deep horizons. Furthermore, C concentrations at depth are relatively low—implying that subsoils have a large capacity to store C and thus might sequester a significant amount of additional atmospheric CO_2_ (Lorenz and Lal, 2005; Minasny et al., 2017; Paustian et al., 2016; Rumpel and Kögel-Knabner, 2011).

A range of processes introduce C to subsoils, including dissolved C transport in percolating water, burial of aboveground litter via physical mixing, and C fluxes from root exudates and root turnover at depth (Rumpel and Kögel-Knabner, 2011). Deep roots in particular have been identified as a potentially useful conduit for increasing subsoil C stocks in managed landscapes (Kell, 2012; Lynch and Wojciechowski, 2015). A large fraction of SOC is root derived, and the depth distribution of SOC correlates with rooting distributions across biomes in natural ecosystems (Grayston et al., 1997; Jobbágy and Jackson, 2000; Rasse et al., 2005). Dead roots and root exudates fuel production of microbial biomass, which subsequently becomes a primary source of mineral-associated C that can persist over long timescales (Sokol et al., 2019). Deeply rooted bioenergy crops can also enhance production of microbial extracellular polysaccharides, cementing soil aggregates that may protect SOC (Sher et al., 2020). In theory, increasing SOC via deep roots might be achieved without displacing conventional food crops if bioenergy crops are grown on marginal lands—which are otherwise not ideal for food production due to low fertility or environmental sensitivity (Gelfand et al., 2013; Lemus and Lal, 2005; Robertson et al., 2017).

While cultivation of perennial bioenergy crops and restoration of perennial grasslands have been widely shown to increase SOC stocks relative to stocks under conventional crops, the majority of studies have focused on the top 30 cm of soil (Anderson-Teixeira et al., 2009; Beniston et al., 2014; Conant et al., 2017; Harris et al., 2015; Monti et al., 2012; Qin et al., 2016). Furthermore, the magnitude of the difference in SOC stocks following conversion to perennial grassland is highly variable (Conant et al., 2017). Predicting the effect of deep roots on subsoil C across different soil types will ultimately require more field studies spanning edaphic gradients that sample deeply (i.e. ≥ 1 m).

Evaluating the effects of deep roots on subsoil C in the field is challenging, however, because differences in SOC stocks between different land use types are often small relative to total SOC stocks (Syswerda et al., 2011). Ideally changes in SOC under different plant types would be quantified in long-term experiments in which initial conditions are controlled and quantified (Liebig et al., 2008; Sanford et al., 2012). An alternative is to sample opportunistically using a paired design (Fisher et al., 1994; Liebig et al., 2005); in this case the plant cover of interest is compared to a neighboring “reference field” representing the conventional management practice and initial conditions are assumed to be the same across the two plots. This approach cannot detect net change SOC over time given that SOC stocks in the reference plot may not be at steady state—but it can detect divergence in SOC stocks under different management scenarios (Sanderman and Baldock, 2010). Furthermore, the paired design can be applied rapidly in locations where initial data are unavailable, enabling wider sampling of edaphic gradients.

Naturally occurring C isotopes (^13^C, ^14^C) can be used as sensitive tracers of C fluxes (Jones and Donnelly, 2004), and are useful for constraining the effect of deep roots on subsoil C when the paired sampling approach is applied (Balesdent et al., 2018; Marin-Spiotta et al., 2009; O’Brien et al., 2013; Richter et al., 1999). For instance, ^13^C is commonly used to quantify the fraction of SOC derived from recent plant inputs in cases where the photosynthetic pathway of the plant cover is replaced, changing the ^13^C signature of the inputs (Balesdent et al., 2018, 1987; Garten and Wullschleger, 2000). However, ^13^C-based mixing models require a clear transition between C_3_ and C_4_ vegetation (Balesdent and Mariotti, 1996), and are thus challenging to apply in agricultural systems with complex cropping histories.

In systems where no clear transition between C_3_ and C_4_ vegetation have occurred, the radioisotope ^14^C provides an alternative to ^13^C. Atmospheric radiocarbon concentrations are sustained by production of ^14^C in the stratosphere, and were elevated by introduction of ^14^C from atomic weapons testing during the 1950’s and 60’s (Hua et al., 2013). Deep soil C exchanges slowly with the atmosphere and thus becomes naturally depleted in ^14^C as it undergoes radioactive decay (Trumbore, 2009). Consequently, recently fixed C introduced to subsoils via increased root production should have an elevated ^14^C signature relative to the preexisting subsoil C pool (Richter et al., 1999). ^14^C can thus provide upper limits on the magnitude of differences in SOC that emerge after replacing conventional crops with deeply rooted crops.

In this paper, we explore C storage in marginal lands cultivated with switchgrass (*Panicum virgatum, L*.), a deeply rooted perennial grass grown as forage and as a cellulosic bioenergy feedstock. We used a paired sampling design at three sites in Oklahoma with different soil textures that experienced soil degradation during the American Dust Bowl and were planted with switchgrass in either 1998 or 2008 and sampled in 2018. Given that 10 years is typically sufficient to measure C stock differences at shallow depths (< 30 cm) when comparing switchgrass to conventional cropland (Anderson-Teixeira et al., 2009), we hypothesized that C stocks at greater depths (> 30 cm) would also diverge between switchgrass and paired reference plots over this timespan. Identifying rates of SOC divergence in subsoils under perennial bioenergy crops is important because the majority of existing studies on land conversion to perennial crops still deal with relatively shallow sampling depths: increasing the number of studies that sample deeply is an imperative for improving regional-to global-scale prediction of perennial crop effects on SOC (Ledo et al., 2020). We tested our hypothesis by quantifying both total C and ^14^C, which we used to develop sensitive estimates of the component of the total C stock that could be attributed to switchgrass.

## 2. Materials and Methods

### 2.1 Field sites

Sampling took place in 2018 at three sites in Oklahoma, USA. At each site, we sampled deep soil cores in >10 year old switchgrass plots and compared these with paired cores collected from nearby fields cultivated with annual crops. The two sites in Southern Oklahoma; Red River farm, Burneyville (hereafter the “Sandy Loam” site; Lat: 33°53’20.52”N, Lon: 97°17’7.13”W) and Pasture Demonstration Farm, Ardmore (hereafter the “Clay” site; Lat: 34°13’11.00”N, Lon: 97°12’36.96”W) had been planted with “Alamo” switchgrass in 2008. The location in Northern Oklahoma, near Stillwater (hereafter the “Loam” site; Lat: 36°8’0.16”N, Lon: 97°6’15.42”W), was planted with “Kanlow” switchgrass in 1998. At the Sandy Loam site, switchgrass was uncut, whereas at the Clay and Loam sites switchgrass was mowed and harvested annually (Loam) or 1-2 times annual (Clay). The switchgrass stands at each site were unfertilized, although the stands at the Clay site were originally established as part of a short-term P response study and thus received fertilizer initially after planting. All three sites were near the outer geographic boundary of the American Dust Bowl during the 1930s and likely experienced wind erosion at that time. Before European settlement, the region likely hosted tall-grass prairie dominated by C_4_ grasses (Cotton et al., 2016). After European settlement in the 19^th^ century, soils in the region were cultivated with C_3_ cereal crops (Paulsen and Shroyer, 2008). The three sites have a broadly similar mean annual climate (Table 1).

**Table 1.**
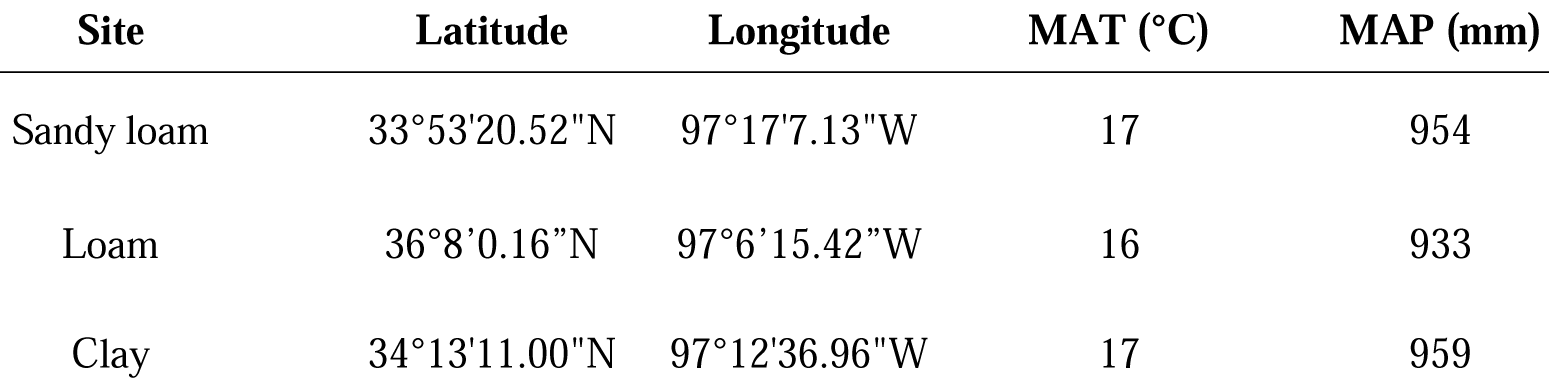
Location and climate of study sites. Mean annual temperature (MAT) and mean annual precipitation (MAP) were obtained using gridded PRISM climate data (Prism Climate Group, Oregon State University, 2011).

| Site | Latitude | Longitude | MAT (°C) | MAP (mm) |
| --- | --- | --- | --- | --- |
| Sandy loam | 33°53'20.52"N | 97°17'7.13"W | 17 | 954 |
| Loam | 36°8'0.16"N | 97°6'15.42"W | 16 | 933 |
| Clay | 34°13'11.00"N | 97°12'36.96"W | 17 | 959 |

At the Sandy Loam site, the reference field had been cultivated with the C_3_ grass rye (*Secale cereal, L*.) in winter and the C_4_ plant crabgrass [*Digitaria sanguinalis, (L.) Scop*.] in the summer for at least the last 15 years under no-till management. Nitrogen fertilizer was applied at approximately 150 kg N ha^-1^ in the reference field annually at this site. At the Clay site, the most recent species grown in the paired reference plots was wheat (*Triticum aestivum, L.)* with a winter cover crop mix; this site was managed with conventional tillage, N was applied at an average rate of 67 kg N ha^-1^ annually, and fields were grazed by cattle in winter. At the Loam site, the reference field was typically planted with wheat and managed with conventional tillage—although during several years throughout 1998-2018 the reference field was planted with the C_4_ grass sorghum [*Sorghum bicolor, (L.) Moench*]; N was applied at an average rate of 72 kg ha^-1^ annually. To our knowledge none of the sites were limed.

The sites spanned a soil texture gradient driven by parent material composition. The Sandy Loam site featured coarse alluvial soils (Coarse-loamy, mixed, superactive, thermic Udic Haplustolls) (National Cooperative Soil Survey, 2020). The Loam site featured soils derived from alluvial and eolian deposits (Fine-loamy, mixed, superactive, thermic Fluventic Haplustolls) (National Cooperative Soil Survey, 2020). Notably, the soils at this site included a buried soil (paleosol) at > 1 m depth. The Clay site included a range of relatively fine-textured soils weathered from Permian shales and sandstones (Fine-loamy, mixed, active, thermic Udic Argiustolls) (National Cooperative Soil Survey, 2020). Soils at the third site varied between clays, clay loams, and sandy clay loams based on the USDA texture classification system; we chose the label “Clay” for this site because it was the most common texture class.

### 2.2 Field sampling

At each site, three soil cores were collected under switchgrass and three cores were collected in an adjacent reference field. We treated the three cores taken in each field as replicate samples, but we acknowledge that these cores are “pseudo-replicated” in that they were collected from the same field, and that a larger sample size would have been ideal (Kravchenko and Robertson, 2011). The low sample size was necessitated by the larger amount of labor required to process > 3 m soil cores and the costs of radiocarbon analyses. Cores were spaced apart in each field so that they would capture within-field variation to the extent possible: cores at the Sandy Loam and the Clay sites were collected in June 2018, from a 20 m^2^ area within each field, and at the Loam site cores were collected Oct 1, 2018, also within a 20 m^2^ area. The reference fields at the Clay and Loam sites were approximately 50 m distant from the switchgrass fields, and the reference field at the Sandy Loam site was approximately 500 m distant but situated in the same soil series. Cores at the Sandy Loam and Cores were taken using a Giddings probe with 10.16 cm (4 inch) inner-diameter tooling and sampling 120 cm intervals. Sampling proceeded to a depth of 3 m unless the probe reached refusal at a shallower depth (this occurred at the Clay site at a depth of 120-150 cm, likely due to calcium cementation at depth). Each core was photographed and divided into 30 cm intervals in the field. The reference plots were chosen to match the soil properties of the switchgrass plots based on field observations.

At all sites, bulk density was estimated by weighing a 4 cm subsample from the center of each core interval in the field and correcting for the gravimetric water content of the subsample to obtain the subsample dry mass. This mass was then divided by the volume of the subsample to calculate bulk density for that interval. Compression during sampling was on-average 7 ± 5% at the Clay site and < 1% at the Loam and Sandy Loam sites. To account for compression during sampling, volumes were linearly corrected over each sampling interval by scaling the observed core length to the expected length (Parfitt et al., 2010). Particles > 2 mm comprised a negligible fraction of the total mass of each interval, and so no correction for rock fraction was performed. C stock calculations were later performed on an equivalent mineral soil mass basis to minimize sensitivity to bulk density estimates (see below).

Roots were removed from the bulk density subsample by hand; approximately 20 person-minutes were spent removing roots per interval. Roots were washed and dried to obtain the root mass in each interval and scaled by the volume of the interval to obtain root biomass estimates. Soil used for total C and C-isotope analysis was sampled from the remainder of the core interval after removing 1 cm from its exterior to exclude soil from upper horizons that might have contaminated the interval during sampling. Soil sampled from the interior of the core was sieved to 2 mm and dried at 105 °C before being subdivided for physical and chemical analysis.

### 2.3 Laboratory analyses

Soil physical and chemical analyses were conducted at the Oregon State University Crop and Soil Science Central Analytical laboratory (https://cropandsoil.oregonstate.edu/cal). Total C and N were quantified by combustion at 1150 °C using an Elementar Macrocube analyzer. Soil texture analysis, soil pH, and exchangeable cations were also quantified on samples from the 0-30, 30-60, and 60-90 cm depth intervals and select intervals at greater depths. Texture was quantified by the sieve and pipette method after removal of organic matter and carbonates (Burt, 2014). Soil pH was measured by electrode in a 1:1 soil:water slurry. Exchangeable cations were quantified by 0.1 M barium chloride extraction and analysis by ICP-OES (Burt, 2014).

Inorganic C was quantified at Lawrence Livermore National Laboratory by treating finely-ground subsamples of each sample with 1 M phosphoric acid in a sealed jar and measuring CO_2_ evolved using a LI-850 infrared gas analyzer (Robertson, 1999). Where carbonates were present, total organic C was obtained by subtracting inorganic C from total C.

C isotopes were quantified on a subset of the soil that was ground to a fine power by hand. Soils that contained carbonates were treated with 1 M HCl to remove inorganic C before isotope analysis. Direct addition of dilute (∼ 1 M) HCl has measurable but relatively small (< 1‰) effects on ^13^C and ^14^C in soils and sediments (Brodie et al., 2011; Komada et al., 2008) and appears to be no more biased than alternative treatment approaches (Brodie et al., 2011). HCl was added to each sample until effervescence ceased and then was allowed to evaporate to prevent leaching of acid-soluble C. Acid-treated soil was analyzed for ^13^C at the University of California Berkeley Center for Stable Isotope Biogeochemistry (https://nature.berkeley.edu/stableisotopelab/). The ^13^C content of each sample (δ^13^C) was reported in per mil (‰) relative to the V-PDB isotope standard. Radiocarbon values were measured on the NEC 1.0 MV Tandem accelerator mass spectrometer (AMS) or the FN Tandem Van de Graaff AMS at the Center for AMS at Lawrence Livermore National Laboratory. Samples were prepared for ^14^C measurement by sealed-tube combustion to CO_2_ in the presence of CuO and Ag and then reduced onto iron powder in the presence of H_2_ (Vogel et al., 1984). The ^14^C content of each sample (Δ^14^C) was reported in ‰ relative to the absolute atmospheric ^14^C activity in 1950. We report Δ^14^C here rather than mean residence times because reporting Δ^14^C does not require an assumption that SOC pools are at equilibrium; negative Δ^14^C values generally indicate less interaction between SOC and the atmosphere and longer SOC residence times. To calculate Δ^14^C, measured δ^13^C values were used to correct for mass dependent fractionation to yield ^14^C activity at a reference δ^13^C of −25‰ (Stuiver and Polach, 1977). Radiocarbon analyses were conducted in late 2018 – 2019 (exact dates are listed for each sample in Supplementary Table 1). Because collection and analysis occurred within a short period, no correction was performed for decay of ^14^C between sampling and analysis. The average instrument uncertainty for Δ^14^C was ± 4 ‰, and the average precision estimated from a set of six duplicate samples was ± 5 ‰.

### 2.4 C stock calculations

We used measured C stocks to directly estimate the net difference in C between the switchgrass and reference fields. We also used ^14^C measurements to develop an indirect estimate that was independent of the measured C stock in the reference field. The C stock calculations were carried out on an equivalent soil mass (ESM) basis using the cumulative coordinate approach (Gifford and Roderick, 2003; Rovira et al., 2015). We used this approach because it is robust to differences in bulk density, and thus better suited to comparing C stocks under different land uses (Wendt and Hauser, 2013). Calculations were performed separately on the surface soil layers—which we defined as the top 500 kg m^-2^ of soil—and the subsoil—which we defined as the 1000 kg m^-2^ of soil directly below the uppermost 500 kg m^-2^ of soil.

We obtained C stocks by using linear interpolation to predict cumulative C mass from cumulative soil mass (Gifford and Roderick, 2003). The mineral mass of each depth interval was used as the basis for developing mass coordinates (Rovira et al., 2015). Mineral mass was obtained by multiplying the mass of the interval by the 1 minus the soil organic matter fraction [soil organic matter fraction = % organic carbon * (1/100) *2; (Pribyl, 2010)]. We then used linear interpolation to develop a piece-wise function defining cumulative OC mass as a function of cumulative mineral soil mass (Gifford and Roderick, 2003):

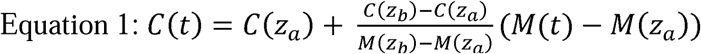

Where C(t) is the cumulative C mass at the target cumulative soil mass M(t), C(z_a_), and C(z_b_) are the cumulative C masses at the upper and lower a boundaries of the sampling interval containing M(t), and M(z_a_) and M(z_b_) are the cumulative mineral masses at those boundaries (Gifford and Roderick, 2003). Using this approach we estimated topsoil C contained in the first 500 kg m^-2^ of soil, and then obtained subsoil C by calculating the total C stock to 1500 kg m^-2^ and subtracting the topsoil C stock. Isotopic values for the topsoil and subsoil were calculated by weighting the values associated with each sampling layer by the contribution of that layer to the C stock. When the lower boundary of the topsoil or subsoil occurred within a layer, isotopic values from that layer were weighted by the C mass that contributed to the topsoil or subsoil.

### 2.5 Isotope calculations

We initially explored the use of ^13^C as a quantitative tracer of switchgrass inputs in our system. The mixed history of C_3_ and C_4_ vegetation at all three sites—and in particular the recent history of periodic C_4_ cropping at the Sandy Loam and Loam sites—suggested that our sites did not experience a clear transition between vegetation types. Depth weighted average δ^13^C values for the subsoil (defined over 500-1500 kg m^-2^ ESM) in the reference plots at our sites ranged between 16.1 and −14.9 ‰, which is at the higher end of the C_4_ plant range (O’Leary, 1988). We measured the δ^13^C of switchgrass roots at the three sites and obtained a range of −13.73 to −13.34 ‰—indicating that the difference between isotopic end-members in a potential ^13^C-based mixing model in the subsoil was only 2-3 ‰. This range is comparable to ∼2 ‰ fractionation effects that apply to plant-tissue end members in isotopic mixing models and are a possible source of uncertainty (Menichetti et al., 2015; Werth and Kuzyakov, 2010). Given these clear limitations, we concluded that δ^13^C—while useful for qualitative interpretation of the SOC depth profiles at our sites—could not be used for identifying switchgrass contributions to SOC quantitatively.

Instead of ^13^C, we used ^14^C to develop estimates of the amount of C introduced to subsoils by switchgrass that were independent of the observed C stocks in the reference plots. The ^14^C signature of plant inputs depends on the composition of the atmosphere, and is thus identical in switchgrass and reference plots. Consequently—while root derived inputs are presumably lower under the reference vegetation—some atmospheric ^14^C is introduced into the subsoil in both cases, and ^14^C can be used to identify net differences in C when comparing the two plots. This contrasts with ^13^C, which is typically used to estimate gross contributions of recently fixed C in the context of paired sampling (Balesdent and Mariotti, 1996).

We did not carry out ^14^C based calculations for the uppermost 500 kg m^-2^ of soil (approximately 30 cm depth) because the Δ^14^C values of the uppermost 500 kg of soil in the reference plots were similar to the range of Δ^14^C value of the recent atmosphere at two of the sites. Specifically, we obtained empirical 95% confidence intervals for the Δ^14^C value of the uppermost 500 kg of soil using Monte-Carlo sampling (see Methods section 2.6 below) spanning [-74, 14] ‰ at the loam site and [-160, −11] ‰ at the Sandy Loam site. These intervals approached or overlapped the Δ^14^C of the recent atmosphere [assumed to be −7 ‰ in 2018 (Hua et al., 2013)], indicating little separation between the isotopic end-members at the surface. This suggests that ^14^C may only be a useful tracer of increased root inputs at depth, where SOC tends to be ^14^C depleted and contrasts strongly with recent inputs.

We divided the subsoil SOC stock under switchgrass (C_s_, kg C m^-2)^ into two parts: (1) a component equal to the C stock under the reference plot (C_r_, kg C m^-2^), representing the initial C stock plus the mass of C equal to what was accrued or lost under the reference vegetation since 1998 or 2008; and (2) a component equal to the additional or “new” C accrued under switchgrass since 1998 or 2008 (C_n_, kg C m^-2^). By definition *C*_*s*_ *= C*_*r*_ *+ C*_*n*_. Each of these components was assigned an accompanying ^14^C signature: Δ_*r*_ and Δ_*s*_, which represented the measured Δ^14^C of the reference and switchgrass plot soils respectively, and Δ_*n*_, which represented the assumed Δ^14^C of *C*_*n*_. These values were related via an isotopic mixing equation:

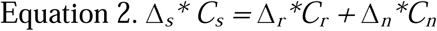

This mixing relationship was used to obtain the fraction (*f*_*n*_) of the C stock under switchgrass comprised by C_n_ and to solve for C_n_:

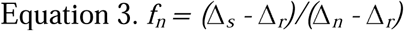

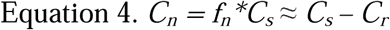

The ^14^C-based isotopic mixing model thus provided an estimate of the C stock difference based on the observed C stock in the switchgrass plot and the shift in ^14^C values between the two plots.

Parameterizing Equation 3 required three Δ^14^C values: Δ_*s*_, Δ_*r*_, and Δ_*n*_. We estimated Δ_s_ and Δ_r_ as the stock-weighted average Δ^14^C values of the subsoils in the switchgrass and reference fields respectively. In contrast, Δ_*n*_ could not be assigned a fixed value because the Δ^14^C of the atmosphere changes over time and there can be lags between root production and integration of root-C into SOC. However, Δ_*n*_ could be constrained within relatively narrow range based on the known atmospheric Δ^14^C and plausible decomposition rates for root-derived SOC since planting. To constrain this range, we modeled the Δ^14^C of SOC produced since 1998 or 2008 using a one-pool soil C model.

The one-pool C model was implemented in SoilR (Sierra et al., 2012) using the function “OnepModel14” and a published atmospheric CO_2_ record for northern hemisphere, extended to 2018 by assuming a 5 ‰ annual decrease in atmospheric Δ^14^C (Hua et al., 2013). The model was initiated in 1998 or 2008 with zero initial C. Inputs were fixed at an arbitrary, constant, nonzero value as the modeled Δ^14^C value was independent of the input rate. While a varying input rate would influence the modeled Δ^14^C value of the SOC, we had no basis for parametrizing a varying rate and the effect of varying inputs was small (e.g. halving litter inputs for the first four years reduced the final Δ^14^C by 4 ‰). The decomposition rate constant was set to two extreme scenarios: either zero (no decomposition) or ln(2) (a one-year half-life). The modelled Δ^14^C value of the SOC pool in 2018 under each scenario was used to define a range for Δ_*n*_. This range spanned from 0 ‰ to +15 ‰ for the Sandy Loam and Clay sites (planted in 2008), and from 0 ‰ to + 44 ‰ for the Loam site (planted in 1998).

### 2.6 Statistical analyses

We evaluated C stock differences between the reference and switchgrass plots by propagating statistical uncertainties using Monte Carlo simulations. Simulations were used to obtain distributions for each estimate of the difference in C stocks between plots given the uncertainties in the input parameters. We obtained 95% confidence intervals from the Monte Carlo distribution of each estimate by computing quantiles of the final distributions (Buckland, 1984), and we obtained empirical p-values from the Monte Carlo intervals to test the hypothesis that the difference in stocks was greater than zero. P-values were obtained using the formula p = (r + 1)/(n + 1), where r was the number of Monte Carlo replicates less than zero and n was the total number of simulations (Davison and Hinkley, 1997). The error in each of the field-measured properties (C stocks and isotope signatures) was modelled by generating normal distributions with the standard deviation and mean obtained from the replicate cores (Huang, 2019). To generate the normal distributions, estimated standard deviations were corrected to account for sample size by dividing them by a correction factor (c_4_) which equals 0.886 when n = 3 (Huang, 2019). The distributions were assumed to vary independently. In the case of Δ_*n*_, we assumed a uniform distribution that ranged between the limiting cases defined in section 2.5 above. Parameter sets were drawn from the distributions 100,000 times. For each parameter set, we calculated one of two quantities: an estimate of *C*_*n*_ from the observed stock difference (*C*_*s*_ *– C*_*r*_) or the ^14^C-based stock difference (*f*_*n*_*\*C*_*s*_).

## 3. Results

### 3.1 Soil physicochemical characteristics

The three sites varied in texture, pH, and exchange properties (Table 2). Clay content and exchangeable cation concentrations were lowest at the Sandy Loam site and highest at the Clay site (Table 2). Ca was the dominant exchangeable cation at the Sandy Loam and Loam sites, whereas Mg and Ca were approximately equal contributors at the Clay site (Table 2). Soil pH values were mildly acid to mildly alkaline across three sites, and exchangeable Al concentrations were below detection, or less than 1% of the total cation pool at all sites, and thus not reported.

**Table 2.** Soil texture and exchange properties. Data are from three replicate cores sampled under switchgrass and paired “reference” annual crops at three sites in Oklahoma characterized by different soil textures. Values represent means of all six cores sampled at each site calculated on averages of the top three depth intervals sampled (0-30, 30-60 and 60-90 cm). Standard deviations are listed in parentheses.

| Site | Particle size (%) |  |  | Exchangeable cations (meq 100g<sup>-1</sup>) |  |  |  | pH |
| --- | --- | --- | --- | --- | --- | --- | --- | --- |
|  | Sand | Silt | Clay | Ca | Mg | Na | K |  |
| Sandy Loam | 63 (3) | 28 (3) | 9 (1) | 3.9 (0.5) | 1.5 (0.2) | N.D | 0.1 (0.04) | 6.2 (0.2) |
| Loam | 41 (6) | 37 (5) | 22 (2) | 8.8 (0.8) | 3.1 (0.4) | N.D | 0.2 (0.03) | 7.1 (0.5) |
| Clay | 46 (10) | 15 (7) | 39 (13) | 7.3 (1.4) | 7.4 (4.0) | 0.8 (1.0) | 0.2 (0.02) | 6.5 (0.4) |

### 3.3 Root biomass

Root biomass values and rooting depth under switchgrass differed substantially between sites. Rooting profiles were deepest at the Sandy Loam site and comparatively shallower at the Loam and Clay sites (Fig 1). Root biomass was much greater under switchgrass at all sites (Fig 1). However, the reference plots were sampled after harvest, and the small number of cores collected (n = 3) may mean that we bypassed roots. Consequently these differences are likely not representative of growing season conditions.

**Figure 1.**
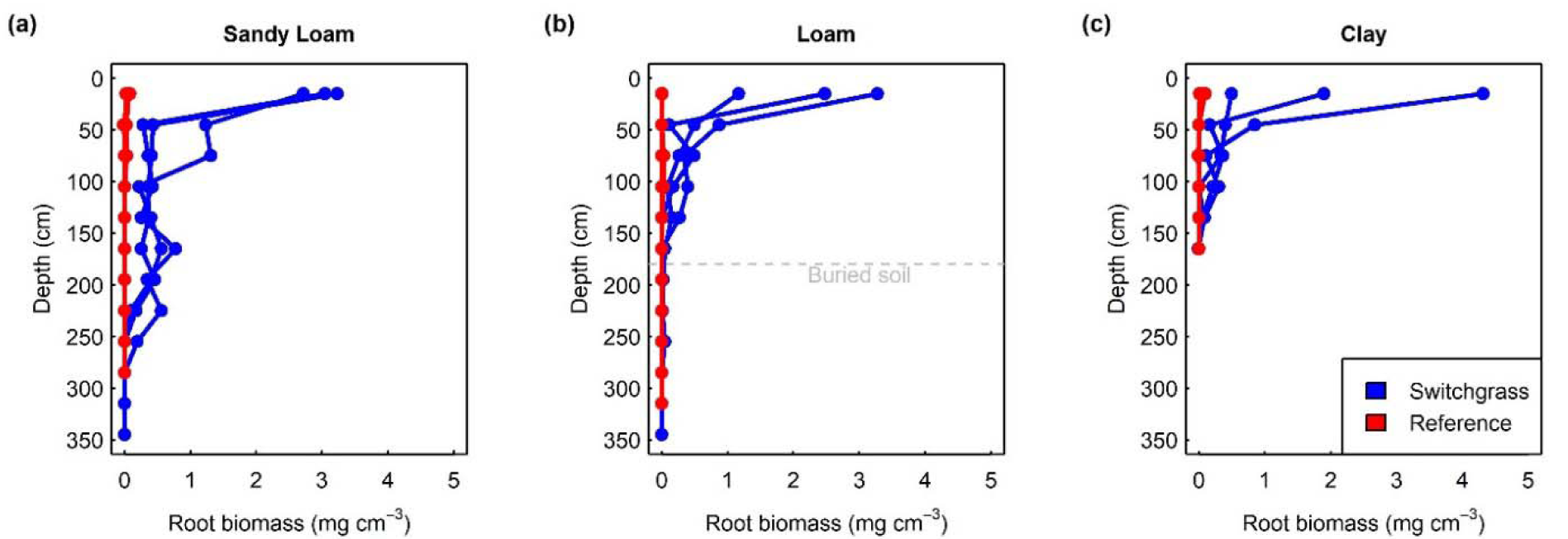
Root biomass versus depth. Data are from three replicate cores sampled under switchgrass and paired “reference” annual crops at three sites in Oklahoma characterized by different soil textures (Sandy Loam, panel a; Loam, panel b; Clay, panel c). Data from each replicate core are shown individually. Cores taken under switchgrass are shown in blue, and cores taken under the reference plot are shown in red. The soil at the Loam site (panel b) featured a buried profile, which is shown with a dashed gray line.

### 3.2 Organic C

Total organic C concentrations were lowest throughout the soil at the Sandy Loam site, intermediate at the Loam site, and highest at the Clay site (Fig 2). At the Sandy Loam site, organic C concentrations were highest in the three cores sampled under switchgrass throughout the uppermost 200 cm of soil (Fig 2a). At the Loam site organic C concentrations were higher in the cores sampled under switchgrass in the top 100 cm of the soil, with the largest difference in the top 30 cm (Fig 2b). We also observed a substantial “bulge” in organic C below 200 cm at the Loam site, which matched the top of the buried paleosol that we identified both in the soil series description and in our field observations. The organic C content of the buried soil was higher in the cores sampled under the reference vegetation (Fig 2b). In contrast to the Sandy Loam and Loam sites, at the Clay site organic C concentrations were generally similar under both vegetation types (Fig 2c).

**Figure 2.**
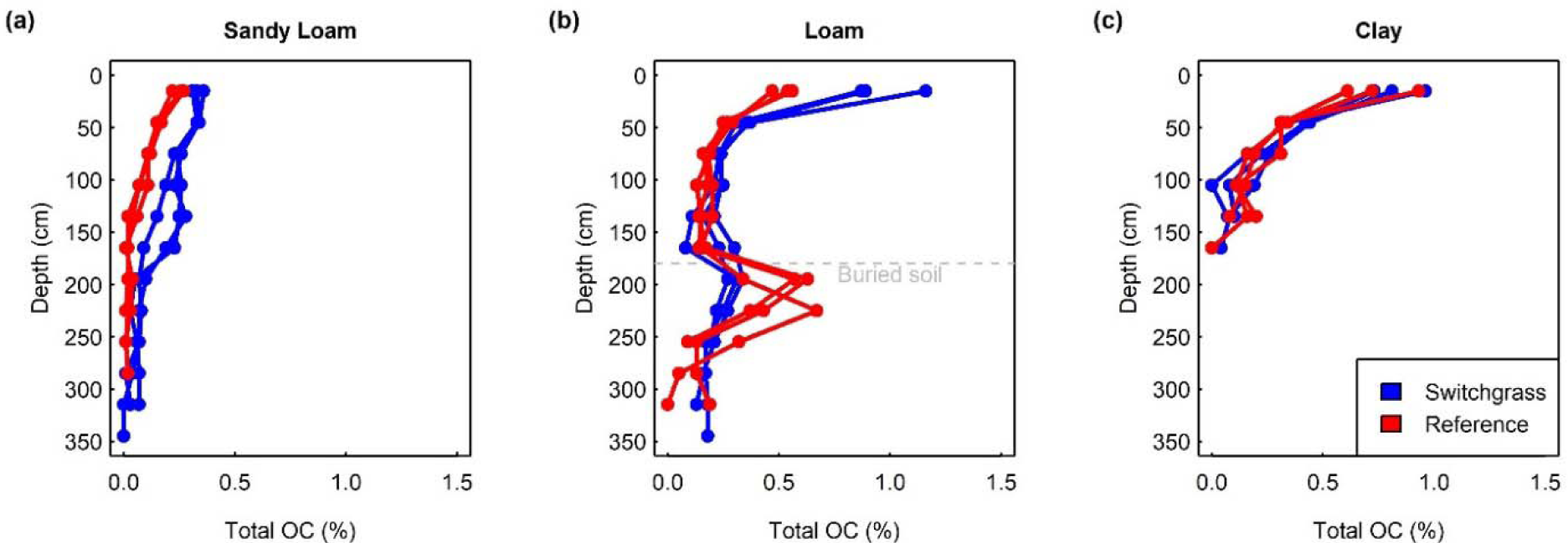
Organic C concentrations versus depth. Data are from three replicate cores sampled under switchgrass and paired “reference” annual crops at three sites in Oklahoma characterized by different soil textures (Sandy Loam, panel a; Loam, panel b; Clay, panel c). Data from each replicate core are shown individually. Cores taken under switchgrass are shown in blue, and cores taken under the reference plot are shown in red. The soil at the Loam site (panel b) featured a buried profile, which is shown with a dashed gray line.

### 3.4 Depth distribution of ^13^C

In general, the ^13^C signature of organic C varied with sampling depth across sites. At the Sandy Loam site, δ^13^C values ranged from −20 to −16 ‰ in the top 30 cm of soil, increased by 3-4 ‰ over 30-90 cm depth, and declined at greater depths (Fig 3a). This pattern appeared under both plant types, but the δ^13^C values were also approximately 2-3 ‰ higher under switchgrass (Fig 3a). At the Loam site, δ^13^C values were also depleted at the surface and comparatively higher at greater depths in a pattern similar to the Sandy Loam site (Fig 3b). The δ^13^C signature was also comparatively higher in cores taken under switchgrass, but this difference attenuated with depth (Fig 3b). At the Clay site, δ^13^C values were highest at the surface and declined with depth (Fig 3c). Patterns under the two plant covers at the Clay site were similar, with slightly higher isotopic values under switchgrass (Fig 3c).

**Figure 3.**
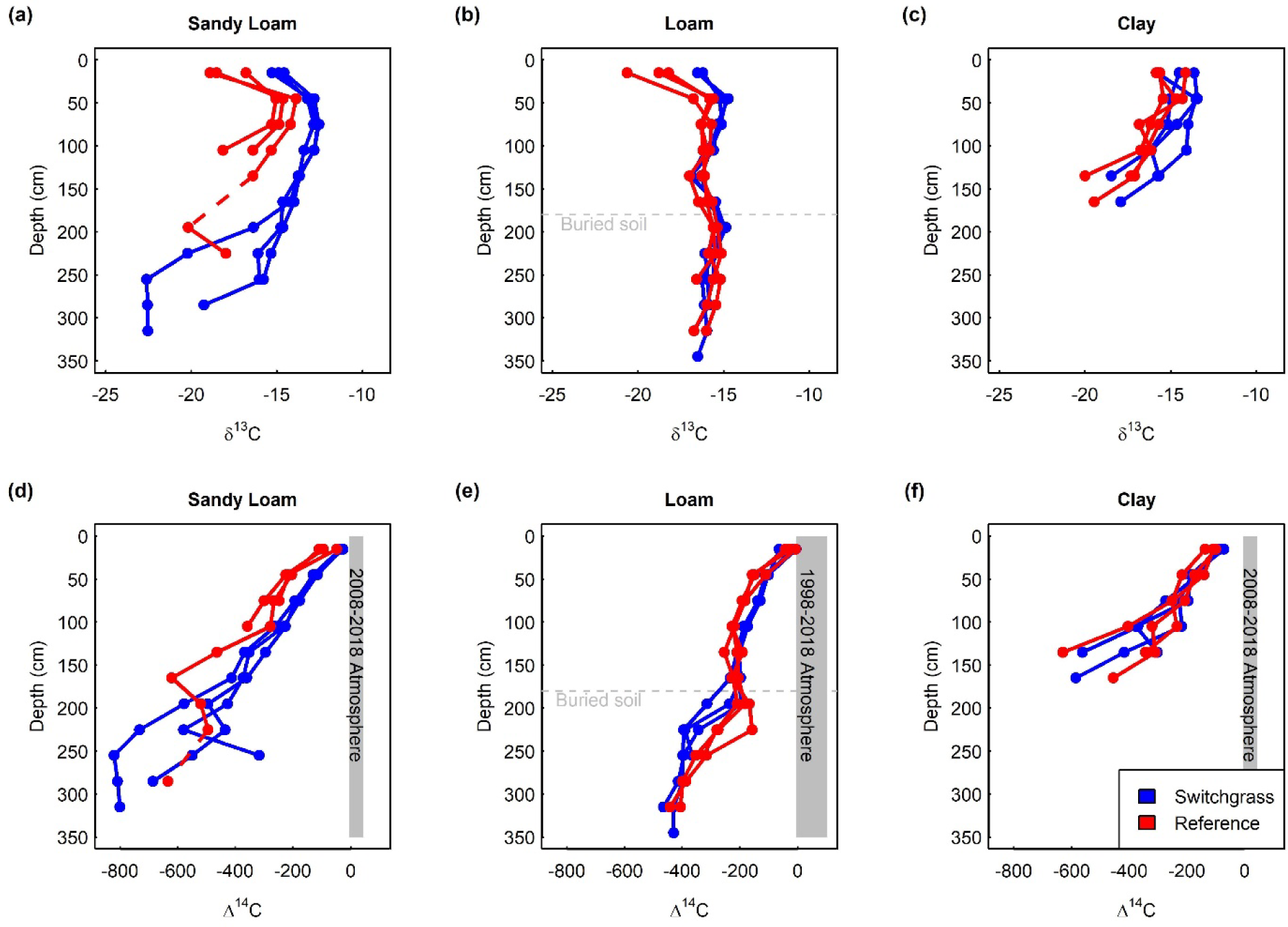
C isotopes versus depth. Data are from three replicate cores sampled under switchgrass and paired “reference” annual crops at three sites in Oklahoma characterized by different soil textures (Sandy Loam, panels a and d; Loam, panels b and e; Clay, panels c and f). Data from each replicate core are shown individually. Cores taken under switchgrass are shown in blue, and cores taken under the reference plot are shown in red. The soil at the Loam site featured a buried profile, which is shown with a dashed gray line. The range of Δ^14^C for the atmosphere over the study period is shown as a gray region on the right of panels d-f. C isotope data could not be collected at all depths at the Sandy Loam site because organic C concentrations were too low; data gaps are interpolated with dashed lines.

### 3.5 Depth distribution of ^14^C

Radiocarbon values declined with depth at all sites (Fig 3d-f). At the Sandy Loam site Δ^14^C values were near zero at the surface and declined to values near −400 ‰ at 150 cm. Below 30 cm, Δ^14^C values were systematically higher in cores taken under switchgrass (Fig 3d). At the Loam site, Δ^14^C values did not decline nearly as steeply as at the Sandy Loam site: at a depth of 150 cm Δ^14^C was approximately 200 ‰. Between 30 and 90 cm the Δ^14^C values of cores sampled under switchgrass were higher at the Loam site (Fig 3e). In the buried soil at the Loam site, Δ^14^C values were higher in cores taken under the reference vegetation (Fig 3e). At the Clay site, Δ^14^C values within the top 30 cm were more depleted relative to the atmosphere than at the other two sites (Fig 3f). The Δ^14^C values declined steeply with depth at the Clay site, reaching values in the −200 to −400 ‰ range at a depth of 1 m. At this site the Δ^14^C depth profiles were broadly similar under the two vegetation types (Fig 3f).

### 3.6 Total organic C stocks

We obtained equivalent soil mass (ESM) estimates of C stocks at each site. The mean C stocks for the top 500 kg m^-2^ of soil (approximately 0-30 cm depth) and the lower 500-1500 kg m^-2^ of soil (approximately 30-100 cm depth) are reported in Table 3. While we focused on ESM estimates when comparing plots to factor out bulk density differences between plots and sites, we also report total estimates to a depth of 1.2 m—which was the greatest depth at which we were able to collect samples across all sites—and to a depth of 2.4 m, which was attained at the Sandy Loam and Loam sites (Table 3). All soil chemical data and C-isotope values are reported in Supplementary Table 1.

**Table 3.** SOC stock estimates. Data are from three replicate cores sampled under switchgrass and paired “reference” annual crops at three sites in Oklahoma characterized by different soil textures. Values are means, with standard deviations in parentheses. The first two columns of data represent stocks estimated on an equivalent soil mass basis; the second two columns represent stocks to a fixed depth. Stocks to 2.4 m are not shown for the Clay site because sampling to this depth was not possible there, possibly due to calcium cementation in the subsoil.

| Site | Plot | Organic C stock (kg m<sup>-2</sup>) |  |  |  |
| --- | --- | --- | --- | --- | --- |
|  |  | 0-500 kg m<sup>-2</sup> | 500-1500 kg m<sup>-2</sup> | 0-1.2 meter | 0-2.4 meter |
| Clay | Switchgrass | 3.9 (0.5) | 2.7 (0.6) | 7.4 (1.0) | n/a |
|  | Reference | 3.6 (0.8) | 2.6 (0.2) | 7.7 (1.1) | n/a |
| Sandy Loam | Switchgrass | 1.7 (0.1) | 2.7 (0.1) | 5.7 (0.3) | 7.3 (0.5) |
|  | Reference | 1.2 (0.1) | 1.3 (0.1) | 2.8 (0.3) | 3.1 (0.3) |
| Loam | Switchgrass | 4.7 (0.8) | 2.8 (0.2) | 8.9 (1.3) | 13.3 (1.4) |
|  | Reference | 2.5 (0.1) | 2.2 (0.1) | 5.9 (0.3) | 11.6 (0.5) |

We compared the C stocks under switchgrass and reference plots (Fig 4). At the Sandy Loam site, direct comparison of the C stocks suggests that there was slightly more C under switchgrass in the top 500 kg m^-2^ of soil (stock difference = 0.4 kg C m^-2^; p < 0.01) and also in the subsoil (stock difference = 1.5 kg C m^-2^; p < 0.01). At the Loam site, we observed significantly more C under switchgrass in the top 500 kg m^-2^ of soil (stock difference = 2.2 kg C m^-2^; p < 0.01) and in the subsoil (stock difference = 0.6 kg C m^-2^; p = 0.01). At the Clay site, the C stock difference in the top 500 kg m^-2^ was comparatively small and not statistically significant (stock difference = 0.2 kg C m^-2^; p = 0.4) and the same was true of the subsoil (stock difference = 0.1 kg C m^-2^; p = 0.44).

**Figure 4.**
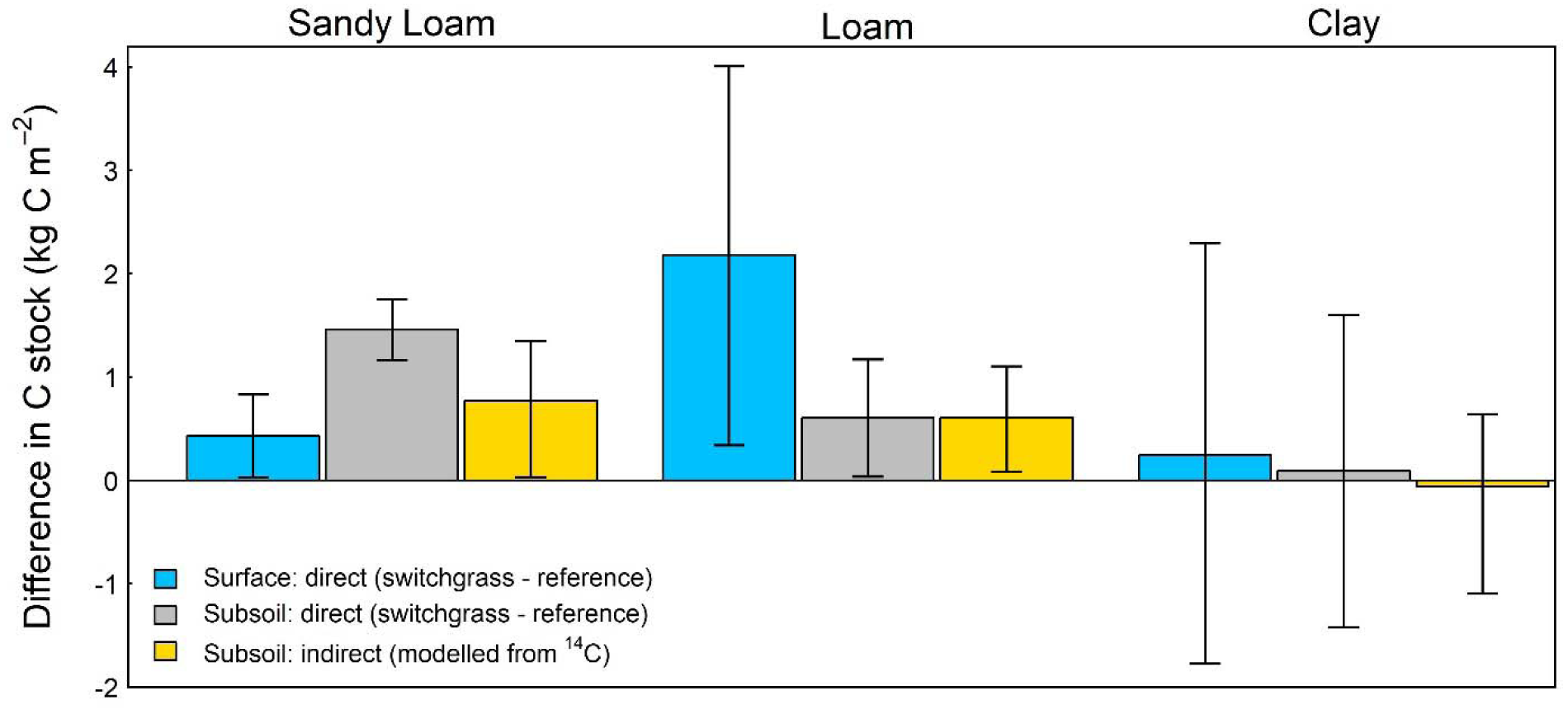
Mean difference in C stock between the switchgrass and annual plant cover. Data are from three replicate cores sampled under switchgrass and paired “reference” annual crops at three sites in Oklahoma characterized by different soil textures. Blue bars show estimates for the top 500 kg m^-2^ of soil, gray bars show estimates for the lower 500-1500 kg m^-2^ of soil, and yellow bars show estimates for the lower 500-1500 kg m^-2^ of soil based on the stock in the switchgrass plot and the shift in Δ^14^C between plots (Equations 2-4). Error bars show 95% confidence intervals derived from Monte Carlo uncertainty propagation.

### 3.7 Stock differences from ^14^C

Using the observed Δ^14^C values, the observed C stocks under switchgrass, and equations 2-4, we developed estimates of the difference in subsoil C stocks between the plots independently of the reference plot C stock (Fig 4). Using Equation 3, we estimated that the fraction of additional C introduced after switchgrass planting (*f*_*n*_) was 0.31 at the Sandy Loam site, 0.21 at the Loam site, and −0.01 (effectively zero) at the Clay site. By multiplying these values by the corresponding C stocks in the switchgrass field we estimated that the ^14^C-based stock difference at the Sandy Loam site was 0.84 kg C m^-2^ (p < 0.01)—which was lower than the direct estimate derived from subtracting the observed C stocks. At the Loam site, the ^14^C-based stock difference was 0.6 kg C m^-2^ (p < 0.01), which overlapped closely with the direct estimate. At the Clay site, the ^14^C-based estimate was near zero and not statistically significant (−0.02 kg C m^-2^; p = 0.48).

## 4. Discussion

### 4.1 Differences in SOC

At two out of the three sites we sampled, we observed significant differences in SOC between switchgrass and reference plots in both topsoil (0-500 kg m^-2^ soil mass, or approximately 0-30 cm) and subsoil (500-1500 kg m^-2^ soil mass, approximately 30-100 cm). At these two sites, differences in subsoil C were in the range of 0.6-1.5 kg C m^-2^. This range is comparable to the value observed in subsoils at 42 paired sites where switchgrass was grown across the upper Midwest (1.2 kg C m^-2^ ; Liebig et al, 2005). If these differences are normalized by the time since planting at each site for the cumulative soil mass of 1500 kg m^-2^ (approximately 0-100 cm depth), the values are 1.9 Mg C ha^-1^ y^-1^ at the Sandy Loam site, 1.4 Mg C ha^-1^ y^-1^ at the Loam site, and 0.3 Mg C ha^-1^ y^-1^ at the Clay site (direct stock comparison). If we use ^14^C-based estimates for the subsoil rather than direct estimates, the time-normalized values are similar: 1.24 Mg C ha^-1^ y^-1^ at the Sandy Loam site, 1.4 Mg C ha^-1^ y^-1^ at the Loam site, and 0.2 Mg C ha^-1^ y^-1^ at the Clay site. These values can be interpreted as “relative changes” in that they estimate the (linear) rate of divergence between switchgrass and conventionally managed systems. This range of rates is typical of switchgrass systems evaluated to a comparable depth (Frank et al., 2004; Qin et al., 2016). Notably, divergence between the two land use types could represent an unknown combination of C sequestration and avoided emissions, depending on the absolute trajectory of C stocks in both fields (Sanderman and Baldock, 2010; Sanford et al., 2012). The discrepancy makes the use of paired plots for C accounting purposes complicated— but at the same time both negative emissions and avoided emissions would be benefits of bioenergy crop production in marginal lands.

### 4.2 Interpreting C isotopes

Both ^13^C and ^14^C were sensitive to land use at the three sites, and in general ^14^C confirmed that larger C stocks under switchgrass at these sites (or lack thereof) can be attributed to recently fixed C in the subsoil. We did, however, discover some disagreement between the directly measured C stocks and the difference estimated using ^14^C: the directly-measured difference in subsoil C stocks was largest at the Sandy Loam, but the shift in ^14^C values at this site was too small to fully accommodate this difference. The simplest interpretation of this result is that the initial C stocks were greater under the switchgrass field before planting—highlighting the limits of the small sample size (n=3) plus the spatially pseudo-replication inherent to the paired sampling design. This interpretation is supported by texture analysis of deeper soil horizons at this site: while soil properties in the upper 90 cm of the soil profile were similar in the reference and switchgrass fields at this site, the reference plot had a higher profile-averaged sand to silt ratio than switchgrass at depths exceeding 90 cm (mean sand/silt = 14 ± 0.7 versus 2 ± 0.4 at a depth of 120 cm, Table S1). This indicates that soil physical characteristics did not match perfectly at this site below a certain depth. At the other two sites where the plots were more closely paired, direct and ^14^C-based methods agreed.

Intriguingly, we observed less total C and comparatively depleted ^14^C values in the buried soil (paleosol) under switchgrass at the Loam site. The ^14^C values in the paleosol were less depleted under the reference plot—and were actually slightly less depleted than the overlying soil (Fig 3e). Given that roots were not observable in the paleosol, we think it is unlikely that patterns in total C and ^14^C at the depth are driven by modern plant cover. Instead, we think it is most likely that the soil under the switchgrass and reference plots—while similar now— experienced different histories, resulting in different C stocks and isotopes at depth. The range of ^14^C values that we observed in the paleosol (−394 to −156 ‰) suggest that it was buried during the mid- or late Holocene (i.e. in the last 5,000 years). It is possible that the paleosol under the switchgrass plot was eroded prior to burial—which would explain its lower C concentrations and ^14^C values relative to the reference plot. The material that was subsequently deposited over both paleosols may have been derived from upland soils containing ^14^C-depleted organic matter, which could explain why the upper part of the paleosol is richer in ^14^C than the overlying base of the modern soil (Lombardo et al., 2018). More generally, deep soil sampling in paired plots can reveal inherited soil features that are not identifiable at the surface—particularly in the mid-continental USA, where paleosols are common under alluvial and eolian deposits (Muhs, 2013).

We generally observed enrichment of ^13^C under switchgrass, particularly at the Sandy Loam site. C_4_ plants like switchgrass have tissue δ^13^C values ranging from −16 to −11 ‰, whereas C_3_ plant tissue ranges from −30 to −20 ‰ (O’Leary, 1988). The shifts we observed are thus consistent with an increase in the abundance of C_4_-derived C under switchgrass. Interpreting the ^13^C data quantitatively is challenging however, given that these sites have experienced a complex history that has included a mix of C_3_ and C_4_ crops. The recent C_3_ plant contribution may explain ^13^C depth profiles at the Sandy Loam and Loam sites, where δ^13^C values in the top 30 cm of soil were lower than those in the subsoil. However, the relatively higher δ^13^C in the subsoil could also reflect fractionation during decomposition (Menichetti et al., 2015; Werth and Kuzyakov, 2010). Given these complexities, it would be challenging to use ^13^C as an unbiased tracer of switchgrass C in the context of our sites—highlighting the value of ^14^C.

### 4.3 Explaining differences between sites

The direct measurements of organic C and the isotopic calculations detected a similar trend: there was more C under switchgrass at the Sandy Loam and Loam sites, and no difference between switchgrass and reference plots at the Clay site. Multiple factors that might explain this pattern given that the three sites have different soil properties and have also experienced different management histories (e.g. tillage, and crop type in the reference fields). Furthermore, the switchgrass stand at the Loam site was 10 years older than the stands at the other two sites. Because these factors are correlated across sites, we have no way to identify which influenced SOC most strongly. Regardless, the large apparent shift at the Sandy Loam site suggests that management effects on C can be substantial even in coarse-textured soils.

## 5. Conclusions

We found that SOC stocks were significantly larger under switchgrass than in nearby reference plots at two out of three sites in Oklahoma. SOC differences were significant at two sites with coarse-textured soils, and not detectable at a site with fine-textured soils. By using ^14^C as a tracer of belowground inputs to the subsoil after planting we were able to confirm that differences in C stocks at the three sites were at least partly attributable to recently fixed C under switchgrass.

This demonstrates that ^14^C can be a useful tracer for divergence of SOC stocks following shifts in cultivation or land use. Further application of ^14^C via repeated measurements and analysis of SOC fractions might help to constrain the trajectory of C stock dynamics, improving C accounting following cultivation of perennial bioenergy crops.

## 6. Acknowledgements

This research is based upon work supported by the U.S. Department of Energy Office of Science, awards SCW1555 and SCW1632 at Lawrence Livermore National Laboratory, award DE-SC0014079 to UC Berkeley and a subcontract to the Noble Research Institute. Lawrence Livermore National Laboratory’s Lab Directed Research and Development program (#19-ERD-010) contributed salary support for ES. Work at LLNL was conducted under the auspices of DOE Contract DE-AC52-07NA27344.

We thank Hugh Aljoe, Kelly Craven, James Rogers and Shawn Norton for facilitating site access and soil collection at the Nobel Research Institute farms. We thank Yanqi Wu for the opportunity to sample Oklahoma State University’s field site near Stillwater, OK. Thanks also to Yuan Wang, Na Ding, Travis Simmons, Josh Barbour, Mellisa McMahon, Jennifer Black, Konstantin Chekhovskiy, Jialiang Kuang, Colin Bates, Ryan Gini, Noah Sokol and Ricardo Feliciano-Rivera for their help with harvests and processing of soil samples.

## References

Anderson-Teixeira, K.J., Davis, S.C., Masters, M.D., Delucia, E.H., 2009. Changes in soil organic carbon under biofuel crops. GCB Bioenergy 1, 75–96. doi: 10.1111/j.1757-1707.2008.01001.x

Balesdent, J., Basile-Doelsch, I., Chadoeuf, J., Cornu, S., Derrien, D., Fekiacova, Z., Hatté, C., 2018. Atmosphere–soil carbon transfer as a function of soil depth. Nature 559, 599–602. doi: 10.1038/s41586-018-0328-3

Balesdent, J., Mariotti, A., 1996. Measurment of turnover of soil organic matter using 13C natural abundance, in: Mass Spectrometry of Soils. Marcel Dekker, New York.

Balesdent, J., Mariotti, A., Guillet, B., 1987. Natural 13C abundance as a tracer for studies of soil organic matter dynamics. Soil Biology and Biochemistry 19, 25–30. doi: 10.1016/0038-0717(87)90120-9

Beniston, J.W., DuPont, S.T., Glover, J.D., Lal, R., Dungait, J.A.J., 2014. Soil organic carbon dynamics 75 years after land-use change in perennial grassland and annual wheat agricultural systems. Biogeochemistry 120, 37–49. doi: 10.1007/s10533-014-9980-3

Brodie, C.R., Leng, M.J., Casford, J.S.L., Kendrick, C.P., Lloyd, J.M., Yongqiang, Z., Bird, M.I., 2011. Evidence for bias in C and N concentrations and δ13C composition of terrestrial and aquatic organic materials due to pre-analysis acid preparation methods. Chemical Geology 282, 67–83. doi: 10.1016/j.chemgeo.2011.01.007

Burt, R., 2014. Soil Survey Laboratory Methods Manual (No 42. Version 5).

Conant, R.T., Cerri, C.E.P., Osborne, B.B., Paustian, K., 2017. Grassland management impacts on soil carbon stocks: a new synthesis. Ecological Applications 27, 662–668. doi: 10.1002/eap.1473

Cotton, J.M., Cerling, T.E., Hoppe, K.A., Mosier, T.M., Still, C.J., 2016. Climate, CO _2_, and the history of North American grasses since the Last Glacial Maximum. Science Advances 2, e1501346. doi: 10.1126/sciadv.1501346

Davison, A.C., Hinkley, D.V., 1997. Bootstrap methods and their application. Cambridge University Press.

Fisher, M.J., Rao, I.M., Ayarza, M.A., Lascano, C.E., Sanz, J.I., Thomas, R.J., Vera, R.R., 1994. Carbon storage by introduced deep-rooted grasses in the South American savannas. Nature 371, 236–238. doi: 10.1038/371236a0

Frank, A.B., Berdahl, J.D., Hanson, J.D., Liebig, M.A., Johnson, H.A., 2004. Biomass and Carbon Partitioning in Switchgrass. Crop Science 44, 1391–1396. doi: 10.2135/cropsci2004.1391

Garten, C.T., Wullschleger, S.D., 2000. Soil Carbon Dynamics beneath Switchgrass as Indicated by Stable Isotope Analysis. Journal of Environmental Quality 29, 645–653. doi: 10.2134/jeq2000.00472425002900020036x

Gelfand, I., Sahajpal, R., Zhang, X., Izaurralde, R.C., Gross, K.L., Robertson, G.P., 2013. Sustainable bioenergy production from marginal lands in the US Midwest. Nature 493, 514–517. doi: 10.1038/nature11811

Gifford, R.M., Roderick, M.L., 2003. Soil carbon stocks and bulk density: spatial or cumulative mass coordinates as a basis of expression? Global Change Biology 9, 1507–1514. doi: 10.1046/j.1365-2486.2003.00677.x

Grayston, S.J., Vaughan, D., Jones, D., 1997. Rhizosphere carbon flow in trees, in comparison with annual plants: the importance of root exudation and its impact on microbial activity and nutrient availability. Applied Soil Ecology 5, 29–56. doi: 10.1016/S0929-1393(96)00126-6

Harris, Z.M., Spake, R., Taylor, G., 2015. Land use change to bioenergy: A meta-analysis of soil carbon and GHG emissions. Biomass and Bioenergy 82, 27–39. doi: 10.1016/j.biombioe.2015.05.008

Harrison, R.B., Footen, P.W., Strahm, B.D., 2010. Deep Soil Horizons: Contribution and Importance to Soil Carbon Pools and in Assessing Whole-Ecosystem Response to Management and Global Change. Forest Science 57, 67–76.

Hua, Q., Barbetti, M., Rakowski, A.Z., 2013. Atmospheric Radiocarbon for the Period 1950– 2010. Radiocarbon 55, 2059–2072. doi: 10.2458/azu_js_rc.v55i2.16177

Huang, H., 2019. Why the scaled and shifted t-distribution should not be used in the Monte Carlo method for estimating measurement uncertainty? Measurement 136, 282–288. doi: 10.1016/j.measurement.2018.12.089

Jobbágy, E.G., Jackson, R.B., 2000. The vertical distribution of soil organic carbon and its relation to climate and vegetation. Ecological Applications 10, 423–436. doi: https://doi.org/10.1890/1051-0761(2000)010[0423:TVDOSO]2.0.CO;2

Jones, M.B., Donnelly, A., 2004. Carbon sequestration in temperate grassland ecosystems and the influence of management, climate and elevated CO2: Tansley review. New Phytologist 164, 423–439. doi: 10.1111/j.1469-8137.2004.01201.x

Kell, D.B., 2012. Large-scale sequestration of atmospheric carbon via plant roots in natural and agricultural ecosystems: why and how. Philosophical Transactions of the Royal Society B: Biological Sciences 367, 1589–1597. doi: 10.1098/rstb.2011.0244

Komada, T., Anderson, M.R., Dorfmeier, C.L., 2008. Carbonate removal from coastal sediments for the determination of organic carbon and its isotopic signatures, δ ^13^ C and Δ ^14^ C: comparison of fumigation and direct acidification by hydrochloric acid: Carbonate removal from coastal sediments. Limnology and Oceanography: Methods 6, 254–262. doi: 10.4319/lom.2008.6.254

Kravchenko, A.N., Robertson, G.P., 2011. Whole-Profile Soil Carbon Stocks: The Danger of Assuming Too Much from Analyses of Too Little. Soil Science Society of America Journal 75, 235–240. doi: 10.2136/sssaj2010.0076

Ledo, A., Smith, P., Zerihun, A., Whitaker, J., VicentelJVicente, J.L., Qin, Z., McNamara, N.P., Zinn, Y.L., Llorente, M., Liebig, M., Kuhnert, M., Dondini, M., Don, A., DiazlJPines, E., Datta, A., Bakka, H., Aguilera, E., Hillier, J., 2020. Changes in soil organic carbon under perennial crops. Global Change Biology gcb.15120. doi: 10.1111/gcb.15120

Lemus, R., Lal, R., 2005. Bioenergy Crops and Carbon Sequestration. Critical Reviews in Plant Sciences 24, 1–21. doi: 10.1080/07352680590910393

Liebig, M.A., Johnson, H.A., Hanson, J.D., Frank, A.B., 2005. Soil carbon under switchgrass stands and cultivated cropland. Biomass and Bioenergy 28, 347–354. doi: 10.1016/j.biombioe.2004.11.004

Liebig, M.A., Schmer, M.R., Vogel, K.P., Mitchell, R.B., 2008. Soil Carbon Storage by Switchgrass Grown for Bioenergy. BioEnergy Research 1, 215–222. doi: 10.1007/s12155-008-9019-5

Lombardo, U., Rodrigues, L., Veit, H., 2018. Alluvial plain dynamics and human occupation in SW Amazonia during the Holocene: A paleosol-based reconstruction. Quaternary Science Reviews 180, 30–41. doi: 10.1016/j.quascirev.2017.11.026

Lorenz, K., Lal, R., 2005. The Depth Distribution of Soil Organic Carbon in Relation to Land Use and Management and the Potential of Carbon Sequestration in Subsoil Horizons, in: Advances in Agronomy. Elsevier, pp. 35–66. doi: 10.1016/S0065-2113(05)88002-2

Lynch, J.P., Wojciechowski, T., 2015. Opportunities and challenges in the subsoil: pathways to deeper rooted crops. Journal of Experimental Botany 66, 2199–2210. doi: 10.1093/jxb/eru508

Marin-Spiotta, E., Silver, W.L., Swanston, C.W., Ostertag, R., 2009. Soil organic matter dynamics during 80 years of reforestation of tropical pastures. Global Change Biology 15, 1584–1597. doi: 10.1111/j.1365-2486.2008.01805.x

Mathieu, J.A., Hatté, C., Balesdent, J., Parent, É., 2015. Deep soil carbon dynamics are driven more by soil type than by climate: a worldwide meta-analysis of radiocarbon profiles. Global Change Biology 21, 4278–4292. doi: 10.1111/gcb.13012

Menichetti, L., Houot, S., van Oort, F., Kätterer, T., Christensen, B.T., Chenu, C., Barré, P., Vasilyeva, N.A., Ekblad, A., 2015. Increase in soil stable carbon isotope ratio relates to loss of organic carbon: results from five long-term bare fallow experiments. Oecologia 177, 811–821. doi: 10.1007/s00442-014-3114-4

Minasny, B., Malone, B.P., McBratney, A.B., Angers, D.A., Arrouays, D., Chambers, A., Chaplot, V., Chen, Z.-S., Cheng, K., Das, B.S., Field, D.J., Gimona, A., Hedley, C.B., Hong, S.Y., Mandal, B., Marchant, B.P., Martin, M., McConkey, B.G., Mulder, V.L., O’Rourke, S., Richer-de-Forges, A.C., Odeh, I., Padarian, J., Paustian, K., Pan, G., Poggio, L., Savin, I., Stolbovoy, V., Stockmann, U., Sulaeman, Y., Tsui, C.-C., Vågen, T.-G., van Wesemael, B., Winowiecki, L., 2017. Soil carbon 4 per mille. Geoderma 292, 59–86. doi: 10.1016/j.geoderma.2017.01.002

Monti, A., Barbanti, L., Zatta, A., Zegada-Lizarazu, W., 2012. The contribution of switchgrass in reducing GHG emissions. GCB Bioenergy 4, 420–434. doi: 10.1111/j.1757-1707.2011.01142.x

Muhs, D.R., 2013. 11.9 Loess and its Geomorphic, Stratigraphic, and Paleoclimatic Significance in the Quaternary, in: Treatise on Geomorphology. Elsevier, pp. 149–183. doi: 10.1016/B978-0-12-374739-6.00302-X

National Cooperative Soil Survey, 2020. National Cooperative Soil Characterization Database.

O’Brien, S.L., Jastrow, J.D., McFarlane, K.J., Guilderson, T.P., Gonzalez-Meler, M.A., 2013. Decadal cycling within long-lived carbon pools revealed by dual isotopic analysis of mineral-associated soil organic matter. Biogeochemistry 112, 111–125. doi: 10.1007/s10533-011-9673-0

O’Leary, M.H., 1988. Carbon Isotopes in Photosynthesis. BioScience 38, 328–336. doi: 10.2307/1310735

Parfitt, R.L., Ross, C., Schipper, L.A., Claydon, J.J., Baisden, W.T., Arnold, G., 2010. Correcting bulk density measurements made with driving hammer equipment. Geoderma 157, 46–50. doi: 10.1016/j.geoderma.2010.03.014

Paulsen, G.M., Shroyer, J.P., 2008. The Early History of Wheat Improvement in the Great Plains. Agronomy Journal 100, S-70-S-78. doi: 10.2134/agronj2006.0355c

Paustian, K., Lehmann, J., Ogle, S., Reay, D., Robertson, G.P., Smith, P., 2016. Climate-smart soils. Nature 532, 49–57. doi: 10.1038/nature17174

Pribyl, D.W., 2010. A critical review of the conventional SOC to SOM conversion factor. Geoderma 156, 75–83. doi: 10.1016/j.geoderma.2010.02.003

Qin, Z., Dunn, J.B., Kwon, H., Mueller, S., Wander, M.M., 2016. Soil carbon sequestration and land use change associated with biofuel production: empirical evidence. GCB Bioenergy 8, 66–80. doi: 10.1111/gcbb.12237

Rasse, D.P., Rumpel, C., Dignac, M.-F., 2005. Is soil carbon mostly root carbon? Mechanisms for a specific stabilisation. Plant and Soil 269, 341–356. doi: 10.1007/s11104-004-0907-y

Richter, D.D., Markewitz, D., Trumbore, S.E., Wells, C.G., 1999. Rapid accumulation and turnover of soil carbon in a re-establishing forest. Nature 400, 56–58. doi: 10.1038/21867

Robertson, G.P. (Ed.), 1999. Standard soil methods for long-term ecological research, Long-term ecological research network series. Oxford University Press, New York.

Robertson, G.P., Hamilton, S.K., Barham, B.L., Dale, B.E., Izaurralde, R.C., Jackson, R.D., Landis, D.A., Swinton, S.M., Thelen, K.D., Tiedje, J.M., 2017. Cellulosic biofuel contributions to a sustainable energy future: Choices and outcomes. Science 356, eaal2324. doi: 10.1126/science.aal2324

Rovira, P., Sauras, T., Salgado, J., Merino, A., 2015. Towards sound comparisons of soil carbon stocks: A proposal based on the cumulative coordinates approach. CATENA 133, 420–431. doi: 10.1016/j.catena.2015.05.020

Rumpel, C., Kögel-Knabner, I., 2011. Deep soil organic matter—a key but poorly understood component of terrestrial C cycle. Plant and Soil 338, 143–158. doi: 10.1007/s11104-010-0391-5

Sanderman, J., Baldock, J.A., 2010. Accounting for soil carbon sequestration in national inventories: a soil scientist’s perspective. Environmental Research Letters 5, 034003. doi: 10.1088/1748-9326/5/3/034003

Sanderman, J., Hengl, T., Fiske, G.J., 2017. Soil carbon debt of 12,000 years of human land use 7.

Sanford, G.R., Posner, J.L., Jackson, R.D., Kucharik, C.J., Hedtcke, J.L., Lin, T.-L., 2012. Soil carbon lost from Mollisols of the North Central U.S.A. with 20 years of agricultural best management practices. Agriculture, Ecosystems & Environment 162, 68–76. doi: 10.1016/j.agee.2012.08.011

Sher, Y., Baker, N.R., Herman, D., Fossum, C., Hale, L., Zhang, X., Nuccio, E., Saha, M., Zhou, J., Pett-Ridge, J., Firestone, M., 2020. Microbial extracellular polysaccharide production and aggregate stability controlled by switchgrass (Panicum virgatum) root biomass and soil water potential. Soil Biology and Biochemistry 143, 107742. doi: 10.1016/j.soilbio.2020.107742

Sierra, C.A., Müller, M., Trumbore, S.E., 2012. Models of soil organic matter decomposition: the SoilR package, version 1.0. Geoscientific Model Development 5, 1045–1060. doi: 10.5194/gmd-5-1045-2012

Sokol, N.W., Kuebbing, Sara.E., Karlsen-Ayala, E., Bradford, M.A., 2019. Evidence for the primacy of living root inputs, not root or shoot litter, in forming soil organic carbon. New Phytologist 221, 233–246. doi: 10.1111/nph.15361

Stuiver, M., Polach, H.A., 1977. Discussion Reporting of ^14^ C Data. Radiocarbon 19, 355–363. doi: 10.1017/S0033822200003672

Syswerda, S.P., Corbin, A.T., Mokma, D.L., Kravchenko, A.N., Robertson, G.P., 2011. Agricultural Management and Soil Carbon Storage in Surface vs. Deep Layers. Soil Science Society of America Journal 75, 92–101. doi: 10.2136/sssaj2009.0414

Trumbore, S., 2009. Radiocarbon and Soil Carbon Dynamics. Annual Review of Earth and Planetary Sciences 37, 47–66. doi: 10.1146/annurev.earth.36.031207.124300

Vogel, J.S., Southon, J.R., Nelson, D.E., Brown, T.A., 1984. Performance of catalytically condensed carbon for use in accelerator mass spectrometry. Nuclear Instruments and Methods in Physics Research Section B: Beam Interactions with Materials and Atoms 5, 289–293. doi: 10.1016/0168-583X(84)90529-9

Wendt, J.W., Hauser, S., 2013. An equivalent soil mass procedure for monitoring soil organic carbon in multiple soil layers. European Journal of Soil Science 64, 58–65. doi: 10.1111/ejss.12002

Werth, M., Kuzyakov, Y., 2010. 13C fractionation at the root–microorganisms–soil interface: A review and outlook for partitioning studies. Soil Biology and Biochemistry 42, 1372–1384. doi: 10.1016/j.soilbio.2010.04.009

